# Computationally efficient demographic history inference from allele frequencies with supervised machine learning

**DOI:** 10.1101/2023.05.24.542158

**Authors:** Linh N. Tran, Connie K. Sun, Travis J. Struck, Mathews Sajan, Ryan N. Gutenkunst

## Abstract

Inferring past demographic history of natural populations from genomic data is of central concern in many studies across research fields. Previously, our group had developed dadi, a widely used demographic history inference method based on the allele frequency spectrum (AFS) and maximum composite likelihood optimization. However, dadi’s optimization procedure can be computationally expensive. Here, we developed donni (demography optimization via neural network inference), a new inference method based on dadi that is more efficient while maintaining comparable inference accuracy. For each dadi-supported demographic model, donni simulates the expected AFS for a range of model parameters then trains a set of Mean Variance Estimation neural networks using the simulated AFS. Trained networks can then be used to instantaneously infer the model parameters from future input data AFS. We demonstrated that for many demographic models, donni can infer some parameters, such as population size changes, very well and other parameters, such as migration rates and times of demographic events, fairly well. Importantly, donni provides both parameter and confidence interval estimates from input AFS with accuracy comparable to parameters inferred by dadi’s likelihood optimization while bypassing its long and computationally intensive evaluation process. donni’s performance demonstrates that supervised machine learning algorithms may be a promising avenue for developing more sustainable and computationally efficient demographic history inference methods.

## INTRODUCTION

Inferring demographic history from genomic data has become routine in many research fields, from elucidating the anthropological origins and migration patterns of modern and archaic human populations (Gutenkunst et al. 2009; Bergström et al. 2020; Marchi et al. 2022; Gopalan et al. 2022), to inferring the population genetic trajectories of endangered animals (Mays Jr et al. 2018; Miller-Butterworth et al. 2021; Chavez et al. 2022). Accounting for demographic history is also essential in setting the appropriate background for detecting signals of natural selection (Nielsen et al. 2005; Boyko et al. 2008; Kim et al. 2017), disease associations (Mathieson & McVean 2012), and recombination hotspots (Johnston & Cutler 2012). Due to the wide range of possible demographic models and high dimensionality of genome sequence data, such analysis often involves computationally expensive modeling. As the size of genomic datasets rapidly grows to thousands of full genomes, there is a great need for more efficient and scalable methods for extracting information from such datasets.

One class of widely used methods infers demographic history from sequence data summarized as an allele frequency spectrum (AFS). An AFS is a multidimensional array with dimension equal to the number of populations being considered in a given demographic model. Each array entry is the number of observed single nucleotide polymorphisms (SNP) with given frequencies in the sampled populations. For example, the [1,2] entry would count SNPs that were singletons in the first population and doubletons in the second. A major advantage of using the AFS as a summary statistic is the ease of scaling to whole genome data (Marchi et al. 2021), as it efficiently reduces the high dimensionality of population genomic data. AFS-based inference methods are, therefore, often fast and suitable for exploring many demographic models (Spence et al. 2018). Given its wide use in countless empirical studies, much progress has been made towards understanding the theoretical guarantees and limitations of the AFS and AFS-based inference (Myers et al. 2008; Achaz 2009; Bhaskar & Song 2014; Terhorst & Song 2015; Baharian & Gravel 2018).

Demographic inference methods based on the AFS often work by maximizing the composite likelihood of the observed AFS under a user-specified demographic history model with parameters such as population sizes, migration rates, and divergence times (Coffman et al. 2016). The expected AFS can be computed via a wide range of approaches (Gutenkunst et al. 2009; Naduvilezhath et al. 2011; Lukić & Hey 2012; Excoffier et al. 2013; Kern & Hey 2017; Jouganous et al. 2017; Kamm et al. 2017) with varying degrees of computational expense, model flexibility, and scalability. Because this is the most computationally intensive step in the procedure, new methods developed thus far have focused on devising algorithms to speed up AFS calculation (Jouganous et al. 2017; Kamm et al. 2017, 2020). However, not much attention has been given to optimizing how the computed AFS is stored and used for inference. In a typical likelihood optimization procedure, hundreds to thousands of expected AFS are computed and compared to the data to obtain the best-fit parameter set. These generated AFS and their corresponding demographic parameters contain information regarding the mapping between the AFS and demographic parameters but are discarded after each optimization run. As there are often a few common demographic models regularly used across studies, if these simulated data could be captured, stored, and distributed for future use, individual groups as well as the research community as a whole could save a lot of time and computational effort by avoiding unnecessary repetition.

The mapping between the AFS and its associated demographic history model parameters can be efficiently captured by supervised machine learning (ML) algorithms. Given a training data set with feature vectors (AFS — input) and labels (demographic history parameters — output), these algorithms can learn the function mapping from the input to the output. While training ML algorithms can be computationally intensive up front, subsequent inference from trained models will have minimal cost (Schrider & Kern 2018). ML algorithms have been widely adopted in population genetics over the past decade, thanks to their efficiency and flexibility. Several studies have used supervised ML algorithms such as random forest (RF) and multilayer perceptron (MLP) with AFS as training data for demographic model selection and demographic parameter inference (Sheehan & Song 2016; Smith et al. 2017; Villanea & Schraiber 2019; Mondal et al. 2019; Lorente-Galdos et al. 2019; Sanchez et al. 2021). In Smith et al. (2017) specifically, the RF algorithm was used to replace the rejection step in the approximate Bayesian computation (ABC) framework, significantly improving overall efficiency (Pudlo et al. 2016). This improvement in efficiency was in part due to more efficient use of simulated data. Whereas in a typical ABC procedure, any simulations beyond a threshold of difference to the data will be discarded, there all simulations were used as input for training the RF classification algorithm. The same principle can be applied in the maximum likelihood optimization and regression framework, where an ML algorithm can be trained by simulated AFS to provide estimates of demographic parameter values, bypassing likelihood optimization.

## NEW APPROACHES

Here, we introduce donni (Demography Optimization via Neural Network Inference), a supervised ML extension to dadi, a widely used AFS-based method for inferring models of demographic history (Gutenkunst et al. 2009) and natural selection (Kim et al. 2017). dadi computes the expected AFS by numerically solving a diffusion approximation to the Wright-Fisher model and uses composite-likelihood maximization to fit the model to the data. While the initial implementation of the software could only handle up to three populations, a recent update supports up to five populations (Gutenkunst 2021). donni uses dadi to generate AFS and demographic parameter labels for training Mean Variance Estimation (MVE) networks (Nix & Weigend 1994) (Fig. 1). Researchers can then use donni’s trained MVE networks to instantaneously infer the parameter values and their associated uncertainty from future AFS input data, obviating the need for likelihood optimization. donni supports a wide range of common demographic parameters that dadi supports, including population sizes, divergence times, continuous migration rates, inbreeding coefficients, and ancestral state misidentification ratios. We show that donni has inference accuracy comparable to dadi but requires less computational resource, even after accounting for the cost of training the MVE networks. Our library of trained networks currently includes all demographic models in the dadi API as well as the models from Portik et al. (2017) pipeline. The supported sample sizes are 10, 20, 40, 80, and 160 haplotypes per population (up to 20 haplotypes only for three-population models). For users who only need to use the trained networks for available demographic models, almost no computation is required. For users who require custom models, we also provide our command-line interface pipeline for generating trained models that can save time compared to running likelihood optimization with dadi. Furthermore, the custom models produced can be contributed to our growing library and shared with the community.

**Figure 1:**
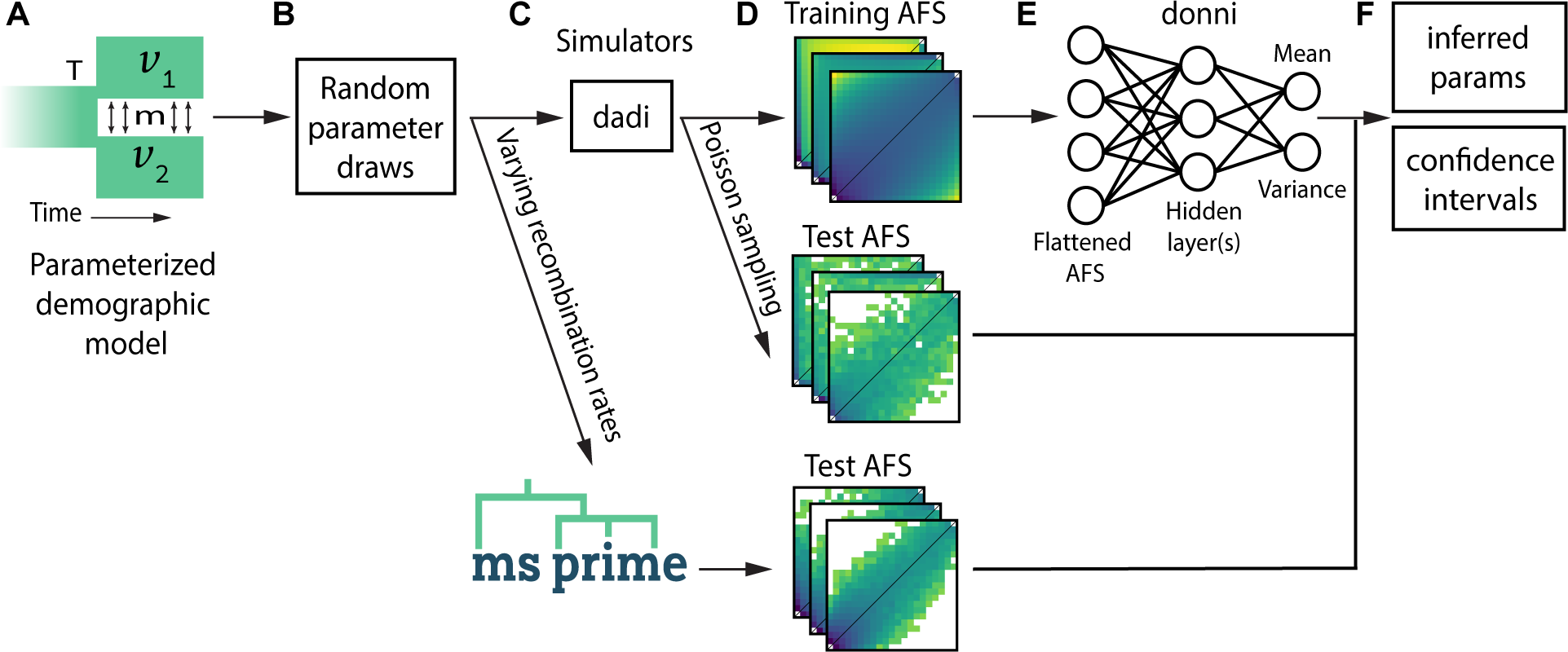
Schematic of the workflow for training and testing donni. For a given demographic model (A), we drew sets of model parameters (B) from a biologically relevant range (Table 1). Each parameter set represents a demographic history and corresponds to an expected AFS. These parameters were input into simulator programs (C) to generate training and test AFS (D). We use the expected AFS simulated with dadi and their corresponding parameters as training data for donni’s MVE networks (E). We generated test data either by Poisson sampling from dadi-simulated AFS or by varying recombination rates with msprime, resulting in a change in variance compared to training AFS. The output of donni’s trained networks includes both inferred parameters and their confidence intervals (F).

## RESULTS

### Choice of MVE network for demographic history model parameter estimation with uncertainty

We wanted to develop a supervised ML method that can infer not only the demographic history parameters but also their associated uncertainties. Uncertainty estimation has not been the focus of previous supervised neural-network-based approaches in demographic history inference (Sheehan & Song 2016; Flagel et al. 2019). There are several techniques for constructing a prediction interval from neural-network-based point estimation as reviewed by (Khosravi et al. 2011). Among them, the MVE method is one of the most conceptually straightforward and least computationally demanding, which are important factors for our goal of improving efficiency.

An MVE network is a feedforward neural network with two output nodes, one for the mean and one for the variance (Fig. 1E). This approach provides an uncertainty estimate in a regression setting by assuming that the errors are normally distributed around the mean estimation. For demographic history inference, the mean is the value of the demographic history model parameter we want to infer. We can construct confidence intervals using the normal distribution defined by the output mean and variance estimates. There are different implementations of the feedforward network architecture for MVE network (Sluijterman et al. 2023). Our implementation is a fully connected network, similar to the MLP, in which all hidden layer weights are shared by the mean and variance output nodes.

### Variance in allele frequencies affects donni training and performance

Since the AFS is the key input data in our method, we first considered how different levels of variance in the AFS might affect training and performance of the MVE networks underlying donni. While the expected AFS computed by dadi under a given set of demographic model parameter gives the mean value of each AFS entry, AFS summarized from observed data will have some variance. We asked whether training the network on AFS with some level of variance or AFS with no variance would lead to better overall performance. When generating AFS simulations, we modeled such variance in the AFS by Poisson-sampling from the expected AFS (examples in Fig. S1A and S2B-D.) We implemented four levels of AFS variance: none, low, moderate, and high in AFS used for training and testing. We then surveyed the inference accuracy for all pairwise combinations for each type of variance in training sets versus test sets.

Overall, we found that networks trained on AFS with no to moderate level of variance perform similarly across all variance levels in test AFS (Fig. S3-S6 for the split-migration model). High variance in training AFS led to substantially poorer performance in parameters that are more difficult to infer, such as time and migration rate. The population size change and ancestral state misidentification parameters were the least affected by AFS variance, and inference accuracy remained similarly high across all variance scenarios. For the time parameter, training on AFS with moderate variance produced the best-performing accuracy across all test cases (Fig. S4). However, for the migration rate parameter, training on AFS with no variance produced the overall best-performing accuracy (Fig. S5). We concluded that for subsequent analyses and model library production for donni, we would train using AFS with no variance, since there was no clear benefit from adding an extra variance simulation step in training. For test AFS, we would use AFS with moderate level of variance to better match real data.

### donni is efficient and has comparable inference accuracy to dadi

Since we built donni to be an alternative to dadi’s likelihood optimization, we compared with dadi in our performance analysis. We validated the inference accuracy of donni for three models: a two-population model with an ancestral population split and symmetric migration between the populations (split-migration model, Fig. 2A), a one-population model with one size change event (two epoch model, Fig. 3A), and a three-population model for human migration out of Africa (the OOA model, Fig. 5A) from Gutenkunst et al. (2009). We also compared the computational efficiency of donni and dadi for two different sample sizes of the split-migration model.

**Figure 2:**
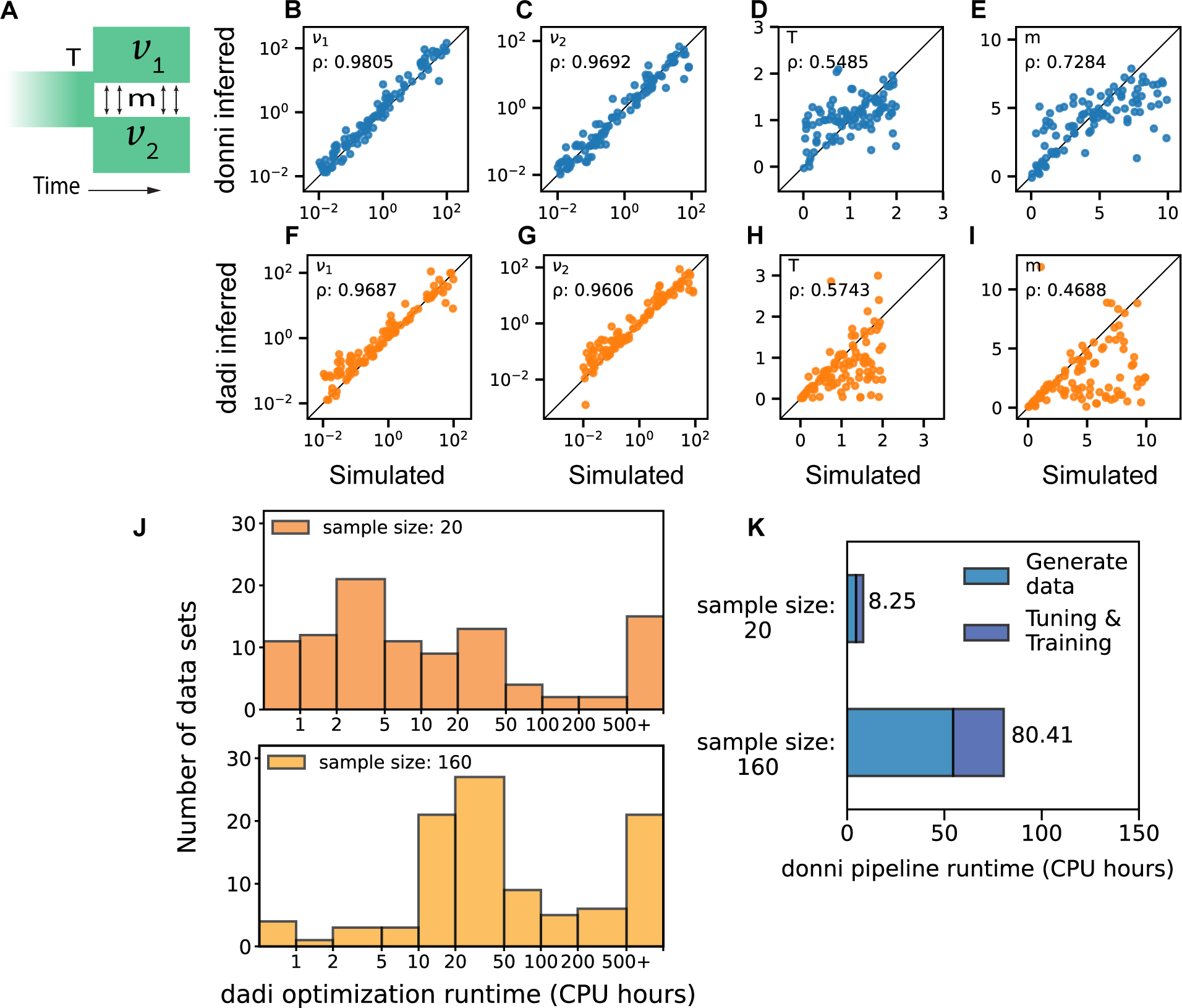
Inference accuracy and computing time of donni and dadi for a two-population model. (A) The two-population split-migration model with four parameters: *ν*_1_ and *ν*_2_ are relative sizes of each population to the ancestral, *T* is time of split, and *m* is the migration rate. (B-I) Inference accuracy by donni (B-E) and dadi (F-I) for the four parameters on 100 test AFS (sample size of 20 haplotypes). (J) Distribution of optimization times among test data sets for dadi. (K) Computing time required for generating donni’s trained networks for two sample sizes. Generate data includes computing time for generating 5000 dadi-simulated AFS as training data. Tuning & training is the total computing time for hyperparameter tuning and training the MVE network using the simulated data.

**Figure 3:**
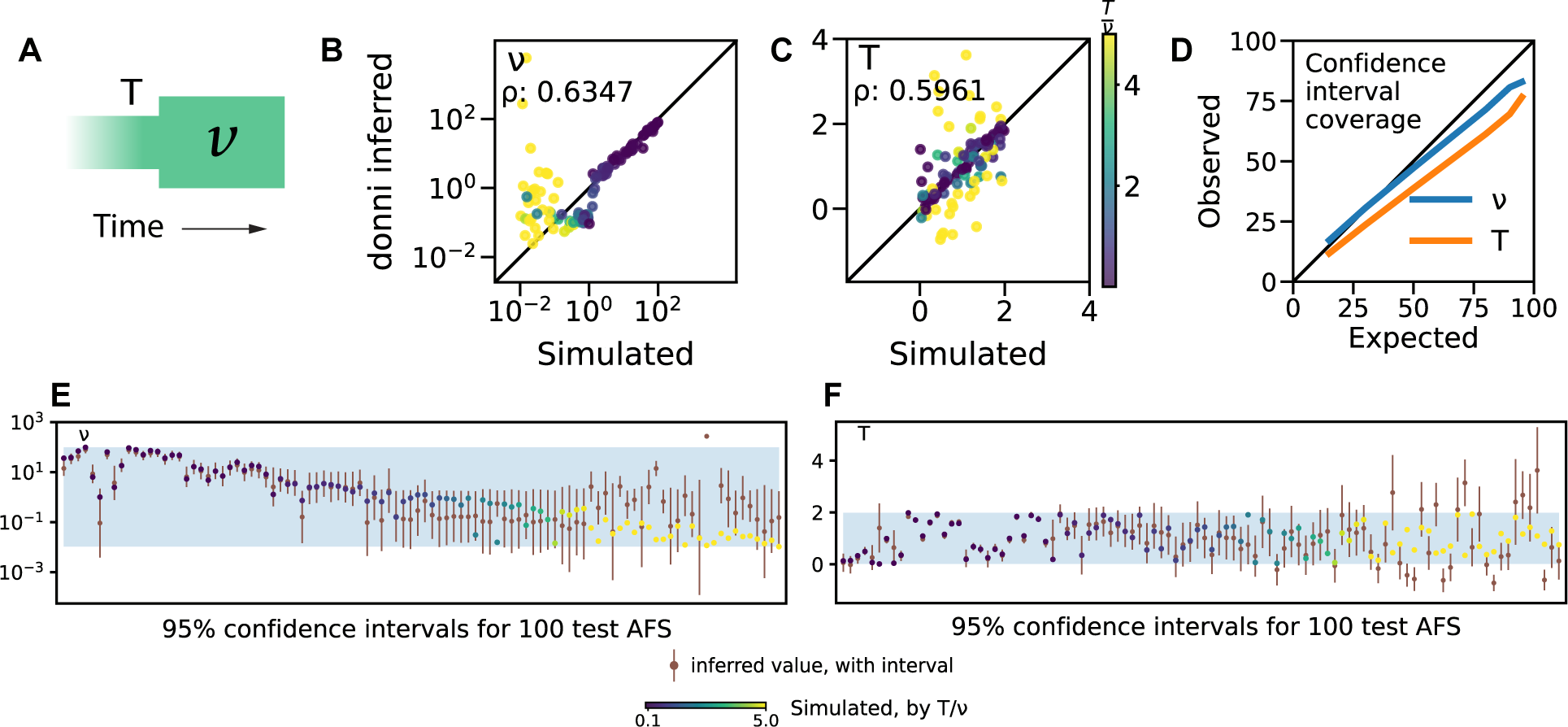
Uninformative AFS affecting inference accuracy and uncertainty quantification method validation. (A) The one-population two-epoch model with two parameters, *ν* for size change and *T* for time of size change. (B-C) Inference accuracy for *ν* and *T* by donni on 100 test AFS, colored by simulated 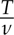 values. (D) Confidence interval coverage for *ν* and *T* by donni. The observed coverage is the percentage of test AFS that have the simulated parameter values captured within the corresponding expected interval. (E-F) As an example, we show details of the 95% confidence interval data points from panel D for 100 test AFS. The simulated values for *ν* (E) and T (F) of these AFS are colored by their 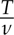 values, similar to panels B-C. donni’s inferred parameter values and 95% confidence interval outputs are in brown. The percentage of simulated color dots lying within donni’s inferred brown interval gives the observed coverage at 95%. The light shades are the simulated parameter range (Table S2) used in simulating training and test AFS. The 100 test AFS are sorted along the x-axis using true 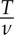 values.

For the split-migration model, donni was able to infer all demographic history parameters with accuracy comparable to dadi (Fig. 2B-I). The population size change parameters *ν*_1_ and *ν*_2_ were inferred very well by both donni (Fig. 2B, C) and dadi (Fig. 2F, G). The time parameter *T* (Fig. 2D, H) and migration rate *m* (Fig. 2E, I) were more difficult to accurately infer for both methods, with dadi having trouble optimizing parameter values close to the specified parameter boundary (Fig. 2E). We used Spearman’s correlation coefficient *ρ* to quantify the monotonic relationship between the true and the inferred parameter values, similar to Flagel et al. (2019). For a more direct measurement of inference accuracy, we also provide the RMSE scores for all models in Table S1.

To compare the efficiency of donni and dadi, we benchmarked the computational resources required by each method to infer demographic parameters from the same 100 test AFS (Fig. 2J-K). Since inferring parameters with donni’s trained networks is computationally trivial, we instead measured the resources required by donni to generate trained networks. For both methods, computation was substantially more expensive as the sample size increased from 20 haplotypes to 160. For dadi (Fig. 2J), there was a spread of optimization runtime among the 100 test AFS, with several difficult spectra requiring more than 500 CPU hours to reach convergence for both sample sizes. By comparison, the computation required for donni (Fig. 2K), including generating training data with dadi, hyperparameter tuning and training, was less than the average time required for running dadi optimization on a single AFS. This result suggests that donni may benefit many cases where dadi optimization can take a long time to reach convergence.

Fig. 2K also suggests that generating the expected AFS with dadi is computationally expensive, often equivalent to if not more so than tuning and training a network. Such expensive operations are indeed what we aimed to minimize with donni. During each dadi optimization, a large number of expected AFS are also calculated. As opposed to discarding all these expensive calculations after each dadi optimization, donni’s trained network effectively captures the mapping between the expected AFS and demographic history model parameter values in its network weights, which can be reused instantaneously in the future.

### donni accurately estimates uncertainty of inferred parameter values

Sometimes, demographic model parameters may be unidentifiable, because multiple parameter sets generate nearly identical AFS. As a simple example, we considered the one-population two epoch model (Fig. 3A), which is parameterized by the relative size *ν* of the contemporary population and the time at which the population size changed *T*. For this model, donni inferences are inaccurate when *T*/*ν* is large (Fig. 3B-C). In this parameter regime, over the time *T* after the size change, the AFS relaxes back to that of an constant-sized equilibrium population. Therefore, in this case, the true parameters are unrecoverable because the AFS itself does not have the appropriate signal to infer them. While this problem may be avoided if users follow the best practice for model selection of exploring simpler models before complex ones (Marchi et al. 2021), it also highlights the need for uncertainty quantification, where a wide confidence interval would appropriately indicate problematic inference.

Using the variance output from the trained MVE networks, donni can calculate any range of confidence intervals specified by the user for each inferred parameter. We validated our uncertainty quantification approach by measuring the observed coverage for six confidence intervals: 15, 30, 50, 75, 80, and 95% intervals (details in Materials & Methods). For the two-epoch model, our approach provided well-calibrated confidence intervals (Fig. 3D). Considering individual test AFS, the uninformative AFS yielded appropriately wide confidence intervals (Fig. 3E-F, yellow points). We found that confidence intervals were similarly well-calibrated for the split-migration model (Fig. S7).

### donni is not biased by linkage between alleles

The Poisson Random Field model underlying dadi (Sawyer & Hartl 1992) and thus donni assumes independence of all genomic loci in the data, which is equivalent to assuming infinite recombination between any pair of loci. But loci within close proximity on the same chromosome are likely sorted together during recombination and therefore linked. To assess how linkage affects donni inference, we tested donni’s networks that were trained on dadi-simulated AFS without linkage on test AFS simulated with msprime, a coalescent simulator that includes linkage (Baumdicker et al. 2022). These msprime-simulated test AFS (examples in Fig. S1B and S2E-G) represent demographic scenarios similar to those in dadi but also include varying levels of linkage under a range of biologically realistic recombination rates. Since smaller recombination rates lead to more linkage and further departure from the training data assumption, we tested donni on AFS with decreasingly small recombination rates down to *r* = 10*^−^*^10^ crossover per base pair per generation, which is two orders of magnitude smaller than the average recombination rate in humans.

Population size parameters *ν* were inferred well no matter the recombination rate, but the inference accuracy for *T* and *m* decreased as the recombination rate decreased (Fig. 4). Confidence intervals were well calibrated at the higher recombination rates (Fig. 4A&E), but too small at the lowest recombination rate (Fig. 4I). These patterns are similar to those we found when testing the effects of AFS variance by Poisson-sampling from expected AFS with dadi (Fig.S3-S7), where accuracy decreased with higher variance, and confidence intervals were underestimated at the highest variances. Note that at *r* = 10*^−^*^10^, linkage disequilibrium often extends entirely across the simulated test regions, so in this regime methods assuming zero recombination, such as IMa3 (Hey et al. 2018), may be more appropriate. Importantly, even though more linkage did lead to higher variance in the estimated parameter values, we did not observe bias in our inferences.

**Figure 4:**
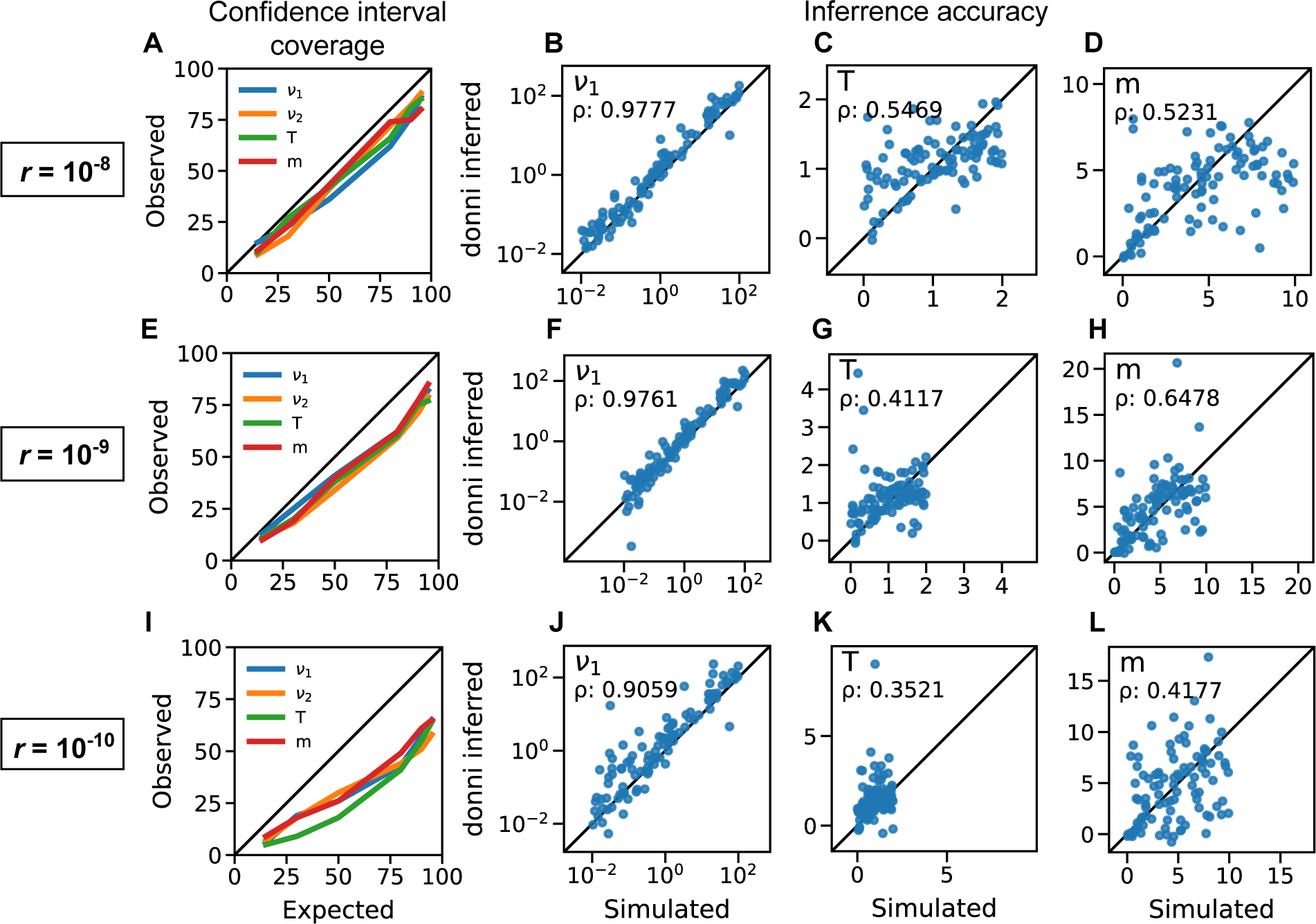
donni’s inference accuracy and uncertainty quantification coverage on msprime-simulated test AFS with linkage. Each row shows the confidence interval coverage and inference accuracy for select parameters of the split-migration demographic model (Fig. 2A) at varying recombination rate. Recombination rate decreases from top to bottom row, corresponding to increased linkage and variance in the msprime-simulated test AFS. The same networks (train on dadi-simulated AFS) were used in this analysis as in Fig. 2F-I.

### Comparison with dadi for the Out-of-Africa model

We tested donni on the three-population Out-of-Africa (OOA) model with 6 size change parameters, 4 migration rates, and 3 time parameters (Fig. 5A). In general, we observed a similar pattern to previous models; size change parameters were often easier to infer than times or migration rates (Fig. 5). For example, both donni and dadi showed near perfect inference accuracy for *ν_A f_* (Fig. 5B&G). They both also performed well for the for *ν_Eu_*, *ν_As_* and *misid* parameters (Fig. S8). But several parameters were challenging for both methods, including some size change parameters, such as *ν_As_*_0_ (Fig. 5C,H), *ν_B_* and *ν_Eu_*_0_ (Fig. S8). The time parameters proved to be the most challenging with relatively lower accuracy for both methods, with *T_A f_* (Fig. 5D,I and *T_B_*S8) being particularly difficult. Overall, both methods agree on the parameters that are easy versus difficult to infer.

**Figure 5:**
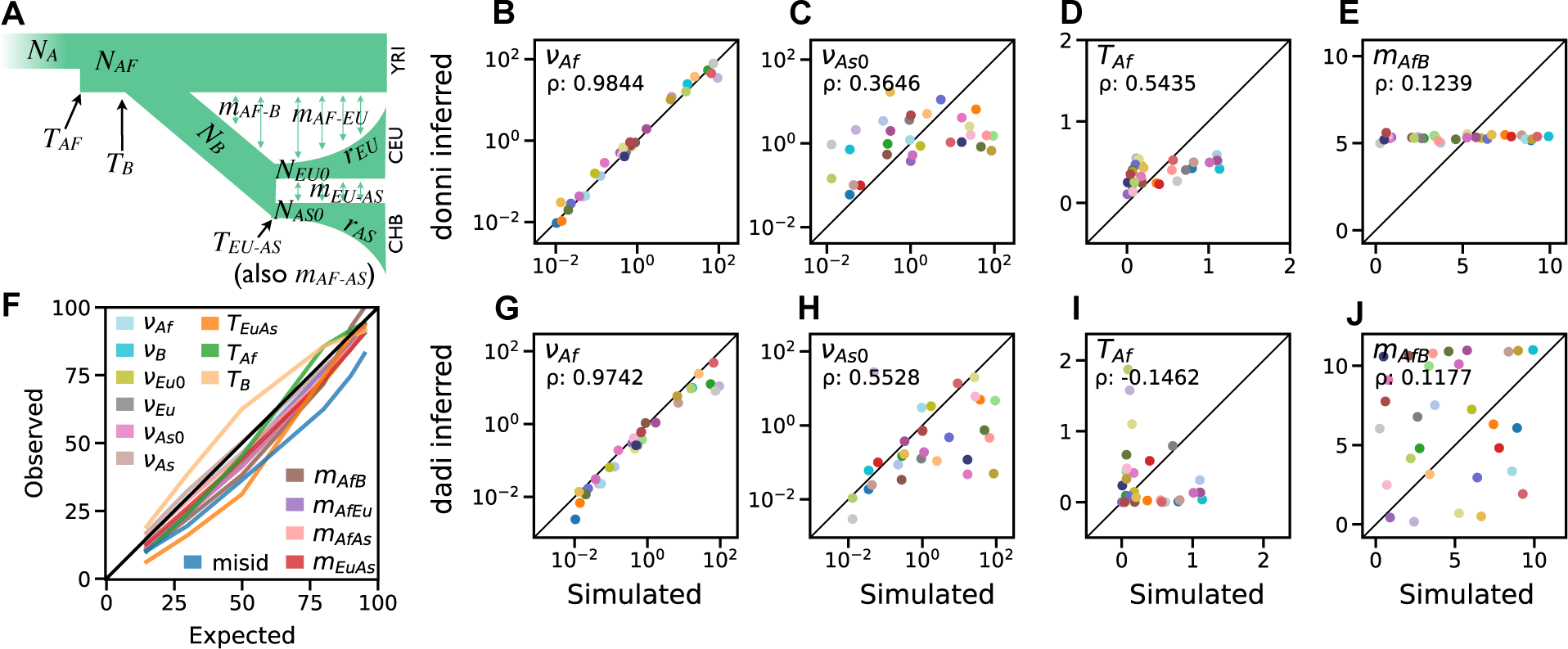
Inference accuracy compared with dadi and confidence interval coverage by donni for the Out-of-Africa demographic model. (A) The three-population Out-of-Africa model with 14 demographic history parameters. (B-E) Inference accuracy for representative parameters on 30 simulated test AFS inferred by donni. (G-J) Inference accuracy for the same parameters and 30 test AFS inferred by dadi. Each of the 30 test AFS is represented by a different color dot. For the accuracy of the rest of the parameters see Fig. S8. (F) donni confidence interval coverage for all model parameters.

However, when inference accuracy is poor on difficult parameters, dadi and donni tend to have different failure patterns. For instance with the *m_A f B_* parameter, dadi tended to get stuck at the parameter boundaries for many AFS (Fig. 5J), while donni essentially inferred the average value for all test AFS (Fig. 5E). This indicates a failure by donni to learn any information from the training AFS for this particular parameter. For all other migration rate parameters in the model, donni performs well, matching dadi (Fig. S8).

While performance varied between the two methods among parameters, donni still had comparable accuracy to dadi in most cases. donni was also able to produce well calibrated confidence intervals for all parameters (Fig. 5F). Due to the computational expense of dadi optimization for this model, we only analyzed 30 test AFS for direct comparison between donni and dadi. Since donni is not as computationally constrained, we also tested donni on all 1000 test AFS per our standard procedure, finding similar results (Table S1).

Finally, we investigated the empirical AFS data from (Gutenkunst et al. 2009) using donni’s trained MVE networks for the Out-of-Africa demographic model (S3). We found that donni’s estimates differ from dadi’s to varying degrees across the parameters. The similarity in accuracy pattern between donni and dadi in Fig. 5 and Fig. S8 does not translate to similar inference values between the two approaches on these data. For example, donni and dadi have similarly high accuracy patterns for *ν_As_* but have very different estimates on the empirical AFS data (*ν_As_* = 7.29 for dadi and *ν_As_* = 1.276 for donni). For this model, donni also tends to infer a stronger migration rate than dadi does, with a higher estimate across all four migration rate parameters. Despite these differences in the estimated parameter values, dadi’s estimates are within donni’s 95% confidence intervals for all parameters.

### donni’s trained networks are accessible

Given its speed, we expect that donni will be useful for quickly exploring many demographic scenarios given a user’s data set. To support this, we have produced trained networks for a large collection demographic history models. These include five one-population and eight two-population models from the current dadi API, plus the 34 two-population and 33 three-population models from Portik et al. (2017). For each of these models, we provide trained networks for unfolded and folded AFS for each of five sample sizes (only two sample sizes for three-population models). For large-scale production, we developed a comprehensive command-line interface pipeline for generating training data, tuning hyperparameters, and assessing the quality of the trained networks. donni’s pipeline is open-source and available on GitHub (https://github.com/lntran26/donni) for users interested in training custom models. The trained network library is available on CyVerse (Merchant et al. 2016; Center 2011) and donni’s command-line interface will automatically download appropriate networks. The library also included all accuracy and confidence interval coverage plots for all supported demographic history models.

## DISCUSSION

We addressed dadi’s computationally intensive optimization procedure by developing donni, a new inference method based on a supervised machine learning algorithm, the MVE network. We found that donni’s trained MVE networks can instantaneously infer many demographic history parameters with accuracy comparable to dadi on simulated data. Even when including computing time required for training the network networks, for many cases donni is faster than dadi’s maximum likelihood optimization. Users are also provided a confidence interval for each inferred demographic history model parameter value from donni. Through examples of one-, two-, and three-population demographic models, we demonstrated that donni’s uncertainty quantification method works well for a wide range of demographic parameters. We also showed that donni works well for AFS simulated by msprime, which includes linkage.

Our approach of using supervised machine learning to reduce the computational expense of the maximum likelihood optimization step is similar in spirit to Smith et al. (2017) using random forests to improve the efficiency of the computationally intensive ABC procedure. While Smith et al. (2017) developed a classification approach for demographic model selection, our method is a regression approach, where we provide a suite of pre-trained regressors for many commonly used demographic history models. Users can quickly explore many possible scenarios and get an estimate for several demographic parameters based on their input AFS data. However, we caution users to always start with simpler models first before trying more complex ones, to avoid exacerbating the uninformative parameter space problem. While we have implemented an accompanying uncertainty quantification tool to aid in identifying such problematic scenarios, best practices in model-based inference should still be followed.

Our choice of AFS as input data for training the network algorithm has several limitations. First, because the size of the AFS depends on the sample size but the network requires a fixed input size, we have to train a different set of networks for different sample sizes within the same demographic history model. Different sets of trained networks are also required for unfolded versus folded AFS. We have limited our trained network library to sample sizes of 10, 20, 40, 80, and 160 haplotypes per population. User data that don’t match exactly these sample sizes will be automatically down-projected (Marth et al. 2004) by donni to the closest available option, leading to some data loss. It is, however, possible to use donni’s pipeline to train custom models that can support a different sample size. We also verified that donni still provides accurate inference and well-calibrated confidence intervals on down-projected data (Fig. S9).

Second, for optimal network performance, we need to normalize the AFS data for training, leading to the loss of information about the parameter *θ* = 4*N_a_µL*, where *N_a_*is the ancestral effective population size, *µ* is the mutation rate and *L* is the sequence length. Estimating *θ* is required for converting all demographic parameters in genetic units to absolute population sizes and divergence times. While donni can provide a point estimate for *θ*, it cannot provide the uncertainty, which is necessary for estimating the uncertainty of absolute parameter values. This limitation can be overcome with a hybrid approach between donni and dadi, where donni’s inferred parameter outputs become the starting point for dadi’s optimization procedure and uncertainty estimation (Coffman et al. 2016). While this approach requires running likelihood optimization, a good starting value provided by donni should reduce overall computing time.

donni trains a separate MVE network for each parameter in a given demographic history model, even though the model parameters are correlated. This is a limitation of our implementation, because the canonical MVE network architecture includes only one node for the mean and one node for the variance. It may be possible to add additional nodes to output means, variances, and covariances from a single network, but we found that this often affects the overall inference quality of the trained MVE network. Additionally, we tested an alternative multi-output regression approach (the scikit-learn Multilayer-Perceptron Regressor) and found that our single-output approach provided similarly accurate estimates (Fig. S10). To our knowledge, existing methods for estimating uncertainties of multi-output neural network regressions are limited.

At its heart, the neural network approach of donni corresponds to a nonlinear regression of model parameters on AFS entries, in contrast to existing approaches which typically maximize a composite likelihood through optimization. Neural networks can be used to estimate likelihoods (e.g., Tejero-Cantero et al. (2020)), which could then be optimized or sampled over, but here we prefer the more direct regression approach. Although dadi and donni display comparable overall accuracy (Fig. 2&5), they may differ when applied to any given data set (Table S3), reflecting differences between regression and composite likelihood optimization.

In conclusion, our results indicate that using supervised machine learning algorithms trained with AFS data is a computationally efficient approach for inferring demographic history from genomic data. Despite implementation limitations discussed above, the AFS is fast to simulate compared to other types of simulated data such as genomic sequence images (Flagel et al. 2019; Sanchez et al. 2021) or coalescent trees (Baumdicker et al. 2022; Kelleher et al. 2016). Furthermore, while ignoring linkage may be a weakness of AFS-based methods, it can also be a strength in that it is more species-agnostic and therefore trained models are transferable among species. A major challenge for AFS-based methods such as ours is the poor scaling to large sample sizes and number of populations, where the AFS matrix becomes high dimensional and sparse, and simulation becomes prohibitively expensive. While we limited this study to three-population models, there have been major improvements in AFS-based methods that can handle more (Gutenkunst 2021; Jouganous et al. 2017; Kamm et al. 2017, 2020). Given our results, a supervised machine learning approach might be a promising next step to extend to such AFS-based methods to further improve their computational efficiency.

## MATERIALS AND METHODS

### Simulations with dadi

We used dadi v.2.3.0 (Gutenkunst et al. 2009) to simulate AFS for training and testing the networks. For each demographic model, we uniformly drew parameter sets from a biologically relevant range of parameters (Table S2). We then generated each expected AFS by specifying the demographic model and parameters in dadi. We calculated the extrapolation grid points used for dadi inte-gration based on the number of haplotypes per population according to Gutenkunst (2021) for one-population models. For models with more than one population, we used the same formula but also increased the grid points by a factor of 1.5 for each additional population. The demographic model parameter values are used as labels for the generated AFS data. To simulate AFS with different levels of variance, we started with the original expected AFS set (no variance). We then Poisson-sampled from the expected AFS to generate a new AFS with variance. We controlled the level of variance by the parameter *θ*, by which we multiplied the expected AFS before sampling. We used *θ* = 10000, 1000, and 100 corresponding to low, moderate, and high levels of variance, respectively (Fig. S3-S7.) Intuitively, modifying *θ* = 4*N_a_µL* is equivalent to altering the effective number of sites surveyed *L*. Assuming *µ ∼* 10*^−^*^8^ and *N_a_ ∼* 10^4^, *θ* = 1000 is equivalent to *L ∼* 2.5 *×* 10^6^ sites. Smaller *θ* is equivalent to fewer sites surveyed, hence noisier AFS. Finally, we normalized both expected and Poisson-sampled AFS for training and testing. The results shown in Fig. 2,3,5, and S8 are based on unfolded AFS with sample size of 20 haplotypes per population.

### Simulations with msprime

We used msprime v1.2.0 (Baumdicker et al. 2022) to simulate AFS from demographic history models while including linkage. We first specified dadi-equivalent demography in msprime for the two epoch and split-migration models. This included the population size change ratio *ν* and time of change *T* parameters for the two epoch model, and population size change ratios *ν*_1_ and *ν*_2_, time *T* of split, and migration rate *m* for the split-migration model. We then specified additional parameters required for msprime to yield *θ* = 4*N_A_Lµ* = 40, 000, with ancestral population size *N_A_* = 10, 000, sequence length *L* = 10^8^ base pairs, and mutation rate *µ* = 10*^−^*^8^ per base pair per generation. We used three recombination rates 10*^−^*^8^, 10*^−^*^9^, and 10*^−^*^10^ per base pair per generation to simulate different levels of linkage and variance in the AFS. We then generated tree-sequence data with msprime before converting to the corresponding unfolded AFS of sample size 20 haplotypes per population and normalizing for testing with trained networks.

### Network architecture and hyperparameter optimization

We used TensorFlow v2.12.1 and Keras v2.12.0 to generate all trained MVE networks for donni. These networks have two fully connected hidden layers containing between 4 and 64 neurons. The exact number of neurons in each hidden layer are hyperparameters that were automatically selected via our tuning procedure described below. The input layer is a flattened AFS with varying sizes depending on the sample size and whether it is a folded or unfolded AFS. The output layer has two nodes for the mean and variance of one demographic history parameter. For tuning and training the network, we implemented a custom loss function based on the negative log-likelihood of a normal distribution:

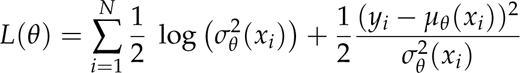

For automatic hyperparameter tuning, we used the HyperBand and RandomSearch tuning algorithms available in keras-tuner v.1.4.6. The 5000 AFS training data set was split 80% for training and 20% for validation. For a given network, we first used HyperBand to optimize both the hidden layer size and learning rate. We then kept the MVE network from HyperBand with the best performance on the validation data, froze the hidden layer size, and then continued tuning only the learning rate using RandomSearch. The MVE network with the best performance on the validation data after RandomSearch is then selected for subsequent training on the full training data. All hyperparameter configurations and non-default settings for the tuning algorithms are listed in Table S4.

### Uncertainty quantification coverage

For uncertainty quantification, the trained MVE network outputs a variance for each inferred demographic history parameter. donni pipeline converts this variance to confidence intervals using the normal distribution. To validate our uncertainty quantification method, we first obtained the method’s estimation for six confidence intervals, 15, 30, 50, 80, 90, and 95% on all test AFS. We then get the observed coverage by calculating the percentage of test AFS that have their corresponding simulated parameter value captured within the estimated interval. The expected versus observed percentages are plotted in our confidence interval coverage plots.

### donni training and testing pipeline

We used 5,000 AFS (no variance) for training and tuning and 1,000 AFS (moderate variance, *θ* = 1000) for accuracy and uncertainty coverage validation. For visualization, only 100 test AFS (30 AFS for the out-of-Africa model) are shown to compare with dadi. However, accuracy scores by donni on all 1000 test AFS are provided in Table S1. Our pipeline tunes and trains one network for each demographic model parameter and sample size. For example, the two epoch model with two parameters *ν* and *T* has 20 independently trained networks: 2 networks for *ν* and *T* times 5 supported sample sizes times 2 polarization states.

### Likelihood optimization with dadi-cli

To infer demographic parameters for a large number of test AFS in parallel (100 AFS for the split-migration model and 30 AFS for the out-of-Africa model), we used dadi’s command-line interface (Huang 2023). We specified the upper and lower bound values for optimization based on the parameter range provided in Table S2. Optimization ran until convergence, as defined by *δlog*(*L*) = 0.0005 for the Out-of-Africa model and *δlog*(*L*) = 0.001 for the split-migration model.

### Benchmarking dadi optimization and donni pipeline

To benchmark the computational expense required for dadi optimization versus for training the networks, we used 10 CPUs on a single computing node for each task. For donni, the tasks are generating training AFS, hyperparameter tuning with HyperBand, and training using the tuned hyperparameters. Estimating demographic parameters for 100 test AFS with donni’s trained networks is nearly instantaneous. For dadi, each test AFS is a task that was optimized until convergence, at which time was recorded, or until the specified cut-off time (50 hours *×* 10 CPUs = 500 CPU hours).

## Acknowledgment

This work was supported by the National Institute of General Medical Sciences of the National Institutes of Health (R01GM127348 and R35GM149235 to R.N.G.).

## Supporting Information

**Figure S1:**
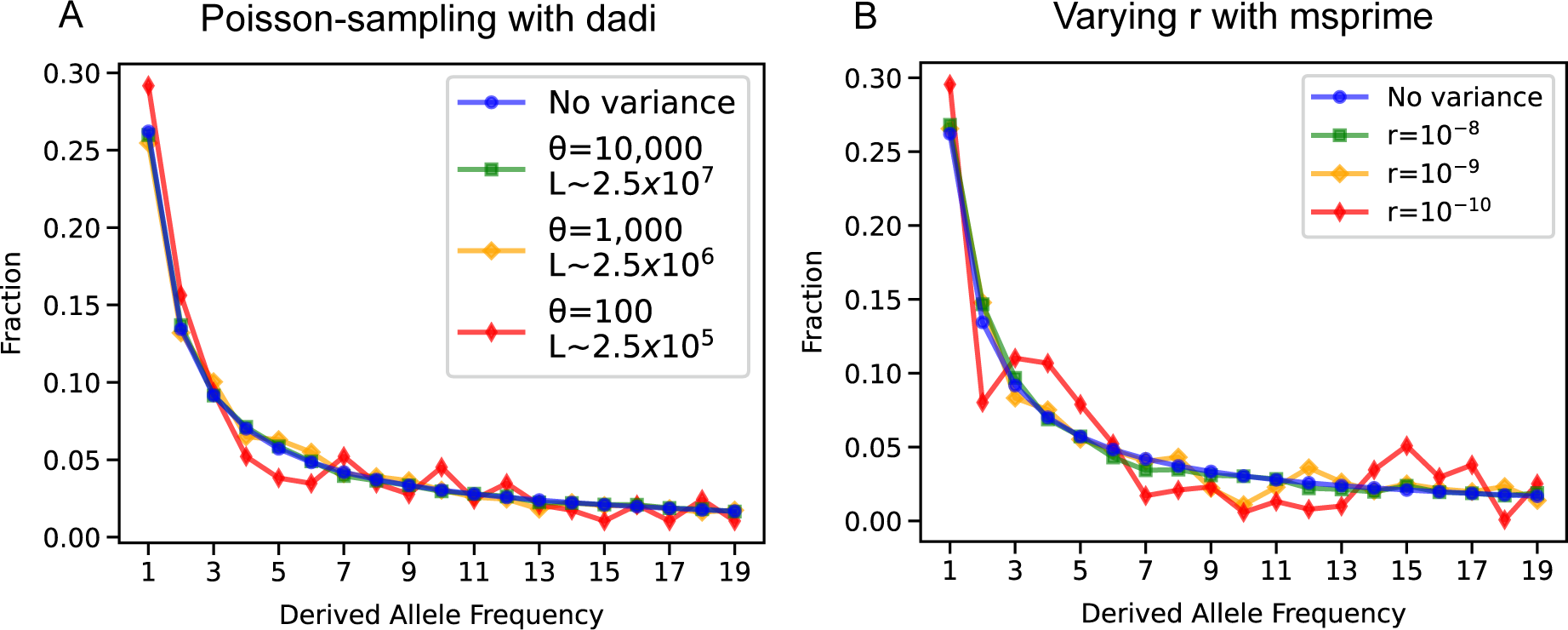
Simulated AFS examples with different variance for the two epoch model. All AFS are normalized and plotted on the same scale. The “No variance” line in both panels is the expected AFS generated by dadi with *ν* = 0.8, *T* = 0.5 (A) AFS with different variance by Poisson-sampling from the “No variance” AFS. (B) msprime-simulated AFS with equivalent demography to (A) but with varying recombination rates. Here *θ* = 4*N_a_µL* = 4 *×* 10^3^ (with *µ* = 10*^−^*^8^ per nucleotide per generation and *L* = 10^8^ base pairs).

**Figure S2:**
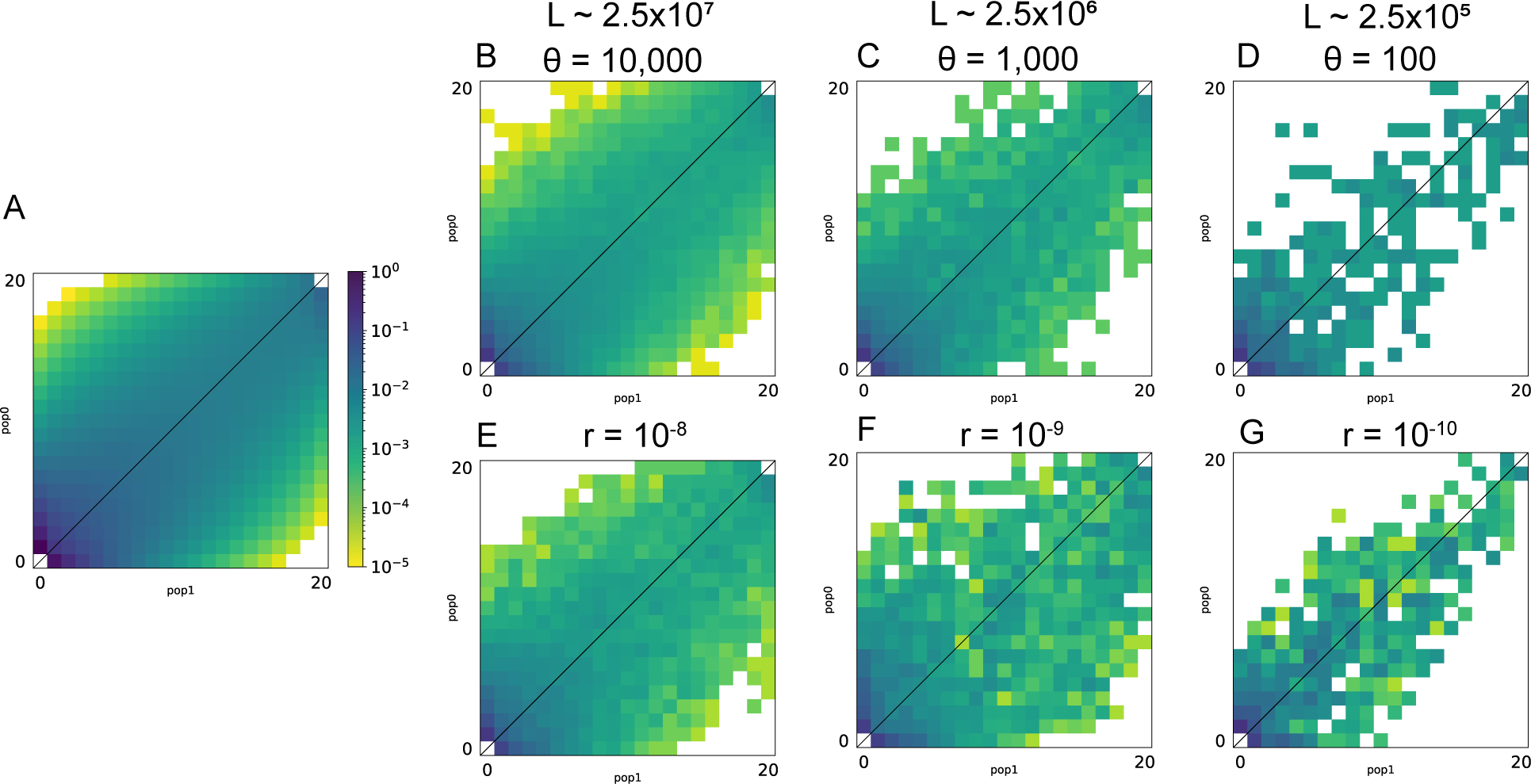
Simulated AFS examples with different variance for the split-migration model. All AFS are normalized and plotted on the same scale. (A) Expected AFS generated by dadi with *ν*_1_ = 1, *ν*_2_ = 0.5, *T* = 2, *m* = 5. (B-D) AFS with different variance by Poisson-sampling from (A). (E-G) msprime-simulated AFS with equivalent demography as in (A) under varying recombination rates.

**Figure S3:**
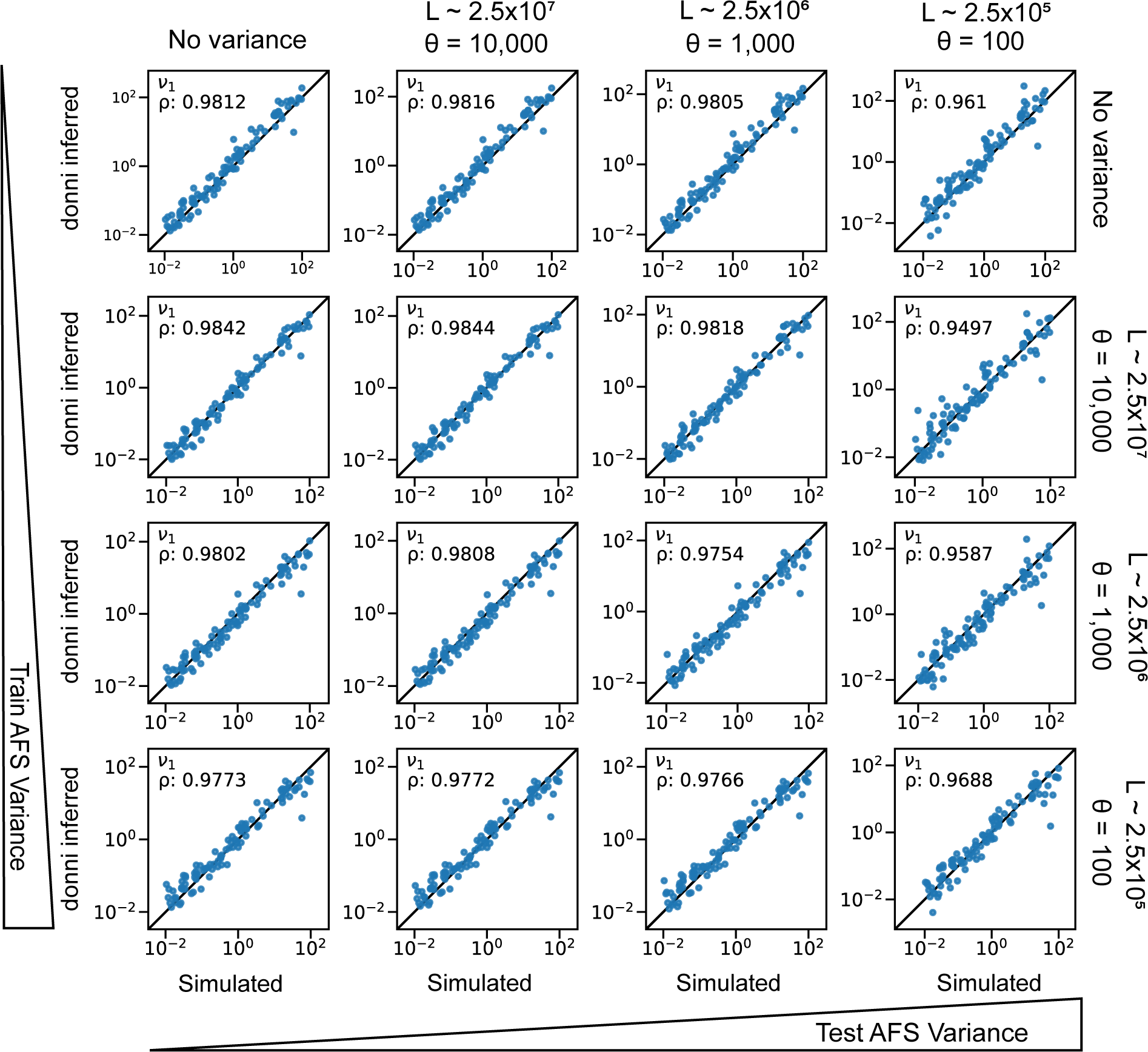
The effects of AFS variance on donni training and performance for the split-migration model size-change parameter. *ν*_1_. Each row corresponds to different levels of variance in training AFS, and each column corresponds to different levels of variance in test AFS. For example, the third panel from the left in the top row is the inference accuracy of a network trained on AFS with no variance tested on AFS with moderate levels of variance (*θ* = 1000 or *L ∼* 2.5 *×* 10^6^ sites surveyed).

**Figure S4:**
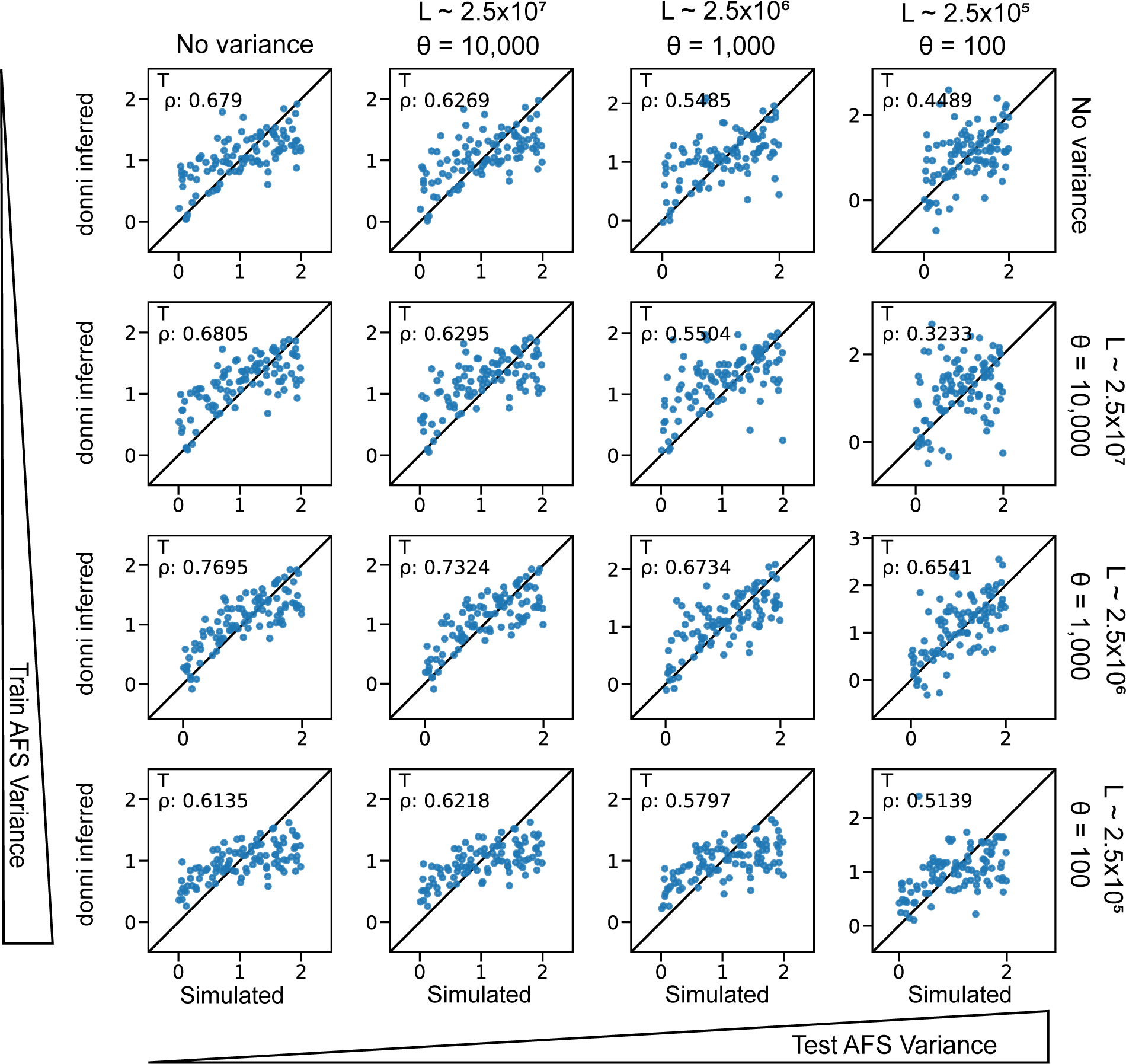
The effects of AFS variance on donni training and performance for the split-migration model time parameter *T*. Panels are as in Fig. S3.

**Figure S5:**
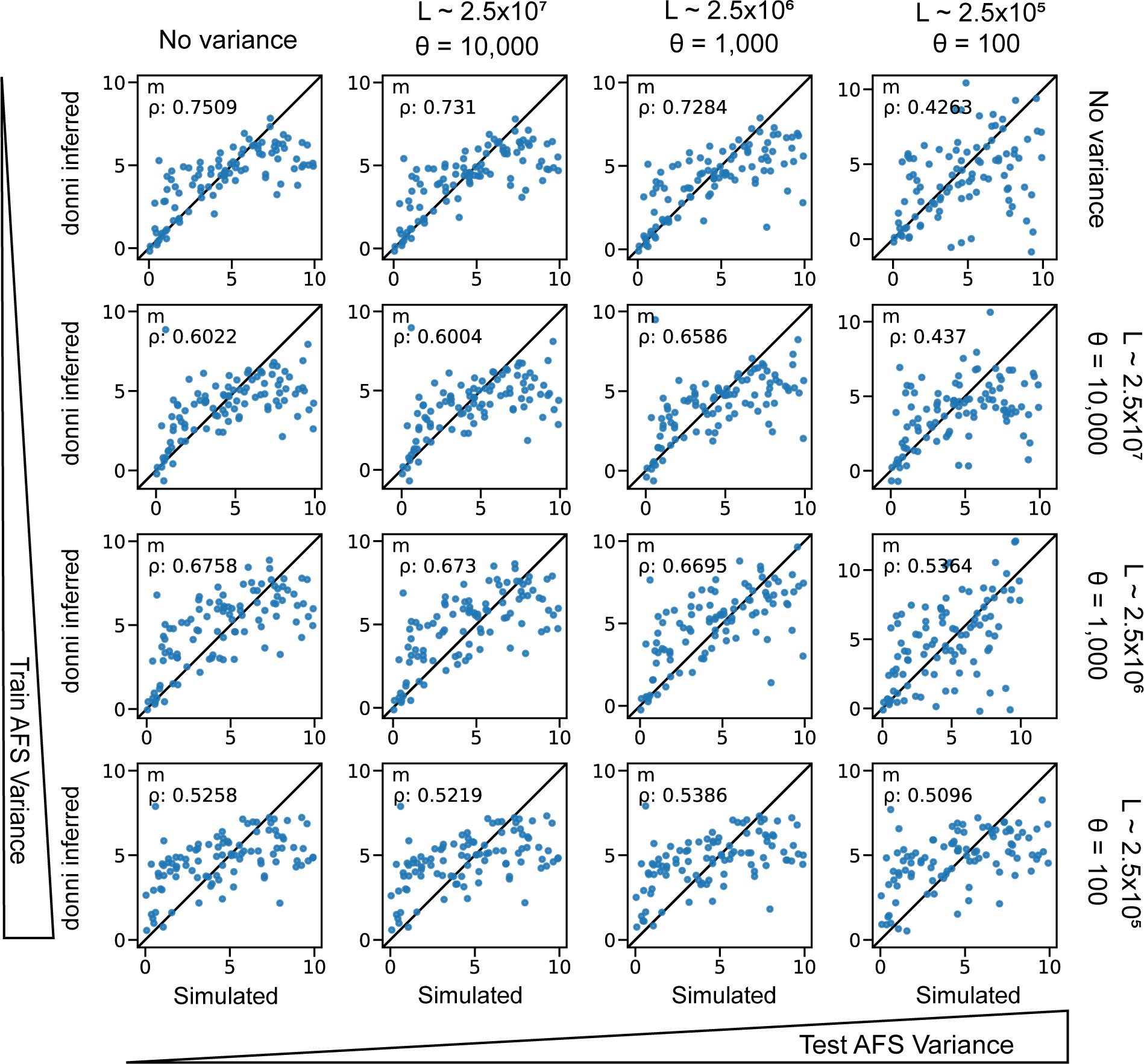
The effects of AFS variance on donni training and performance for the split-migration model migration rate parameter *m*. Panels are as in Fig. S3.

**Figure S6:**
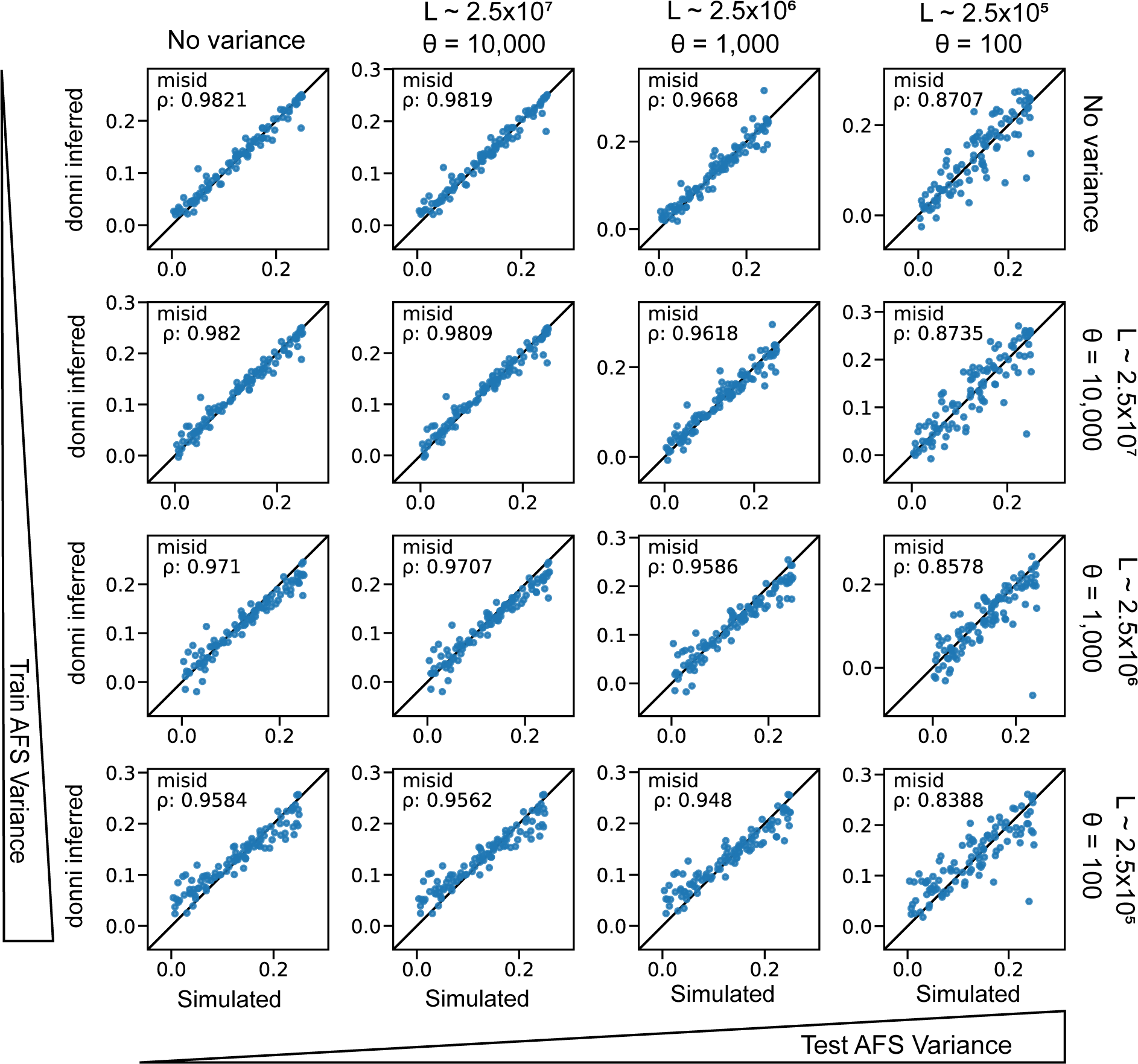
The effects of AFS variance on donni training and performance for the split-migration model ancestral state misidentification parameter. Panels are as in Fig. S3.

**Figure S7:**
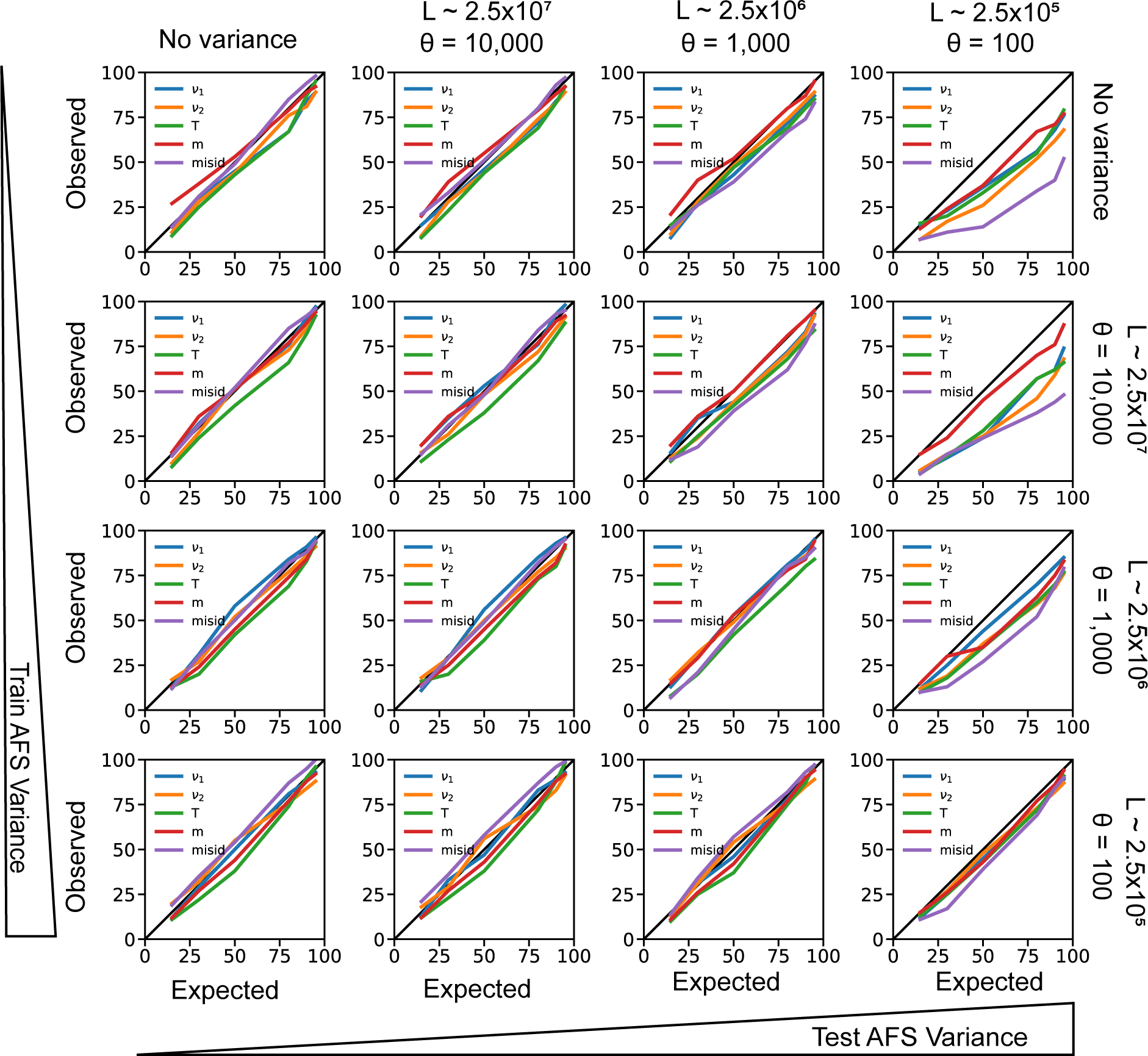
The effects of AFS variance on donni’s uncertainty quantification method for the split-migration model. Panels are as in Fig. S3.

**Figure S8:**
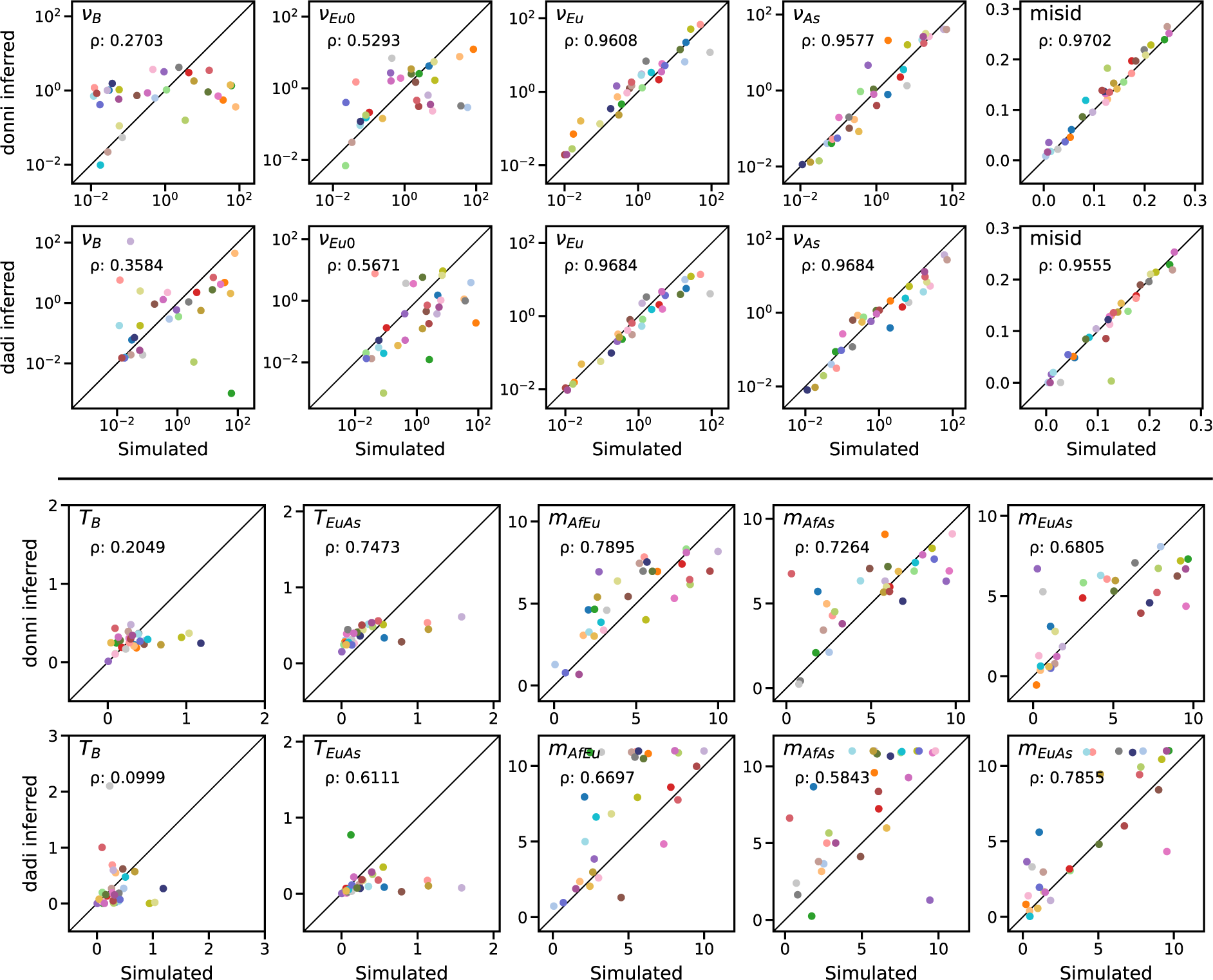
Inference accuracy of dadi and donni on the rest of Out-of-Africa model parameters. Each of the 30 test AFS is represented by a different color dot as in Fig. 5.

**Figure S9:**
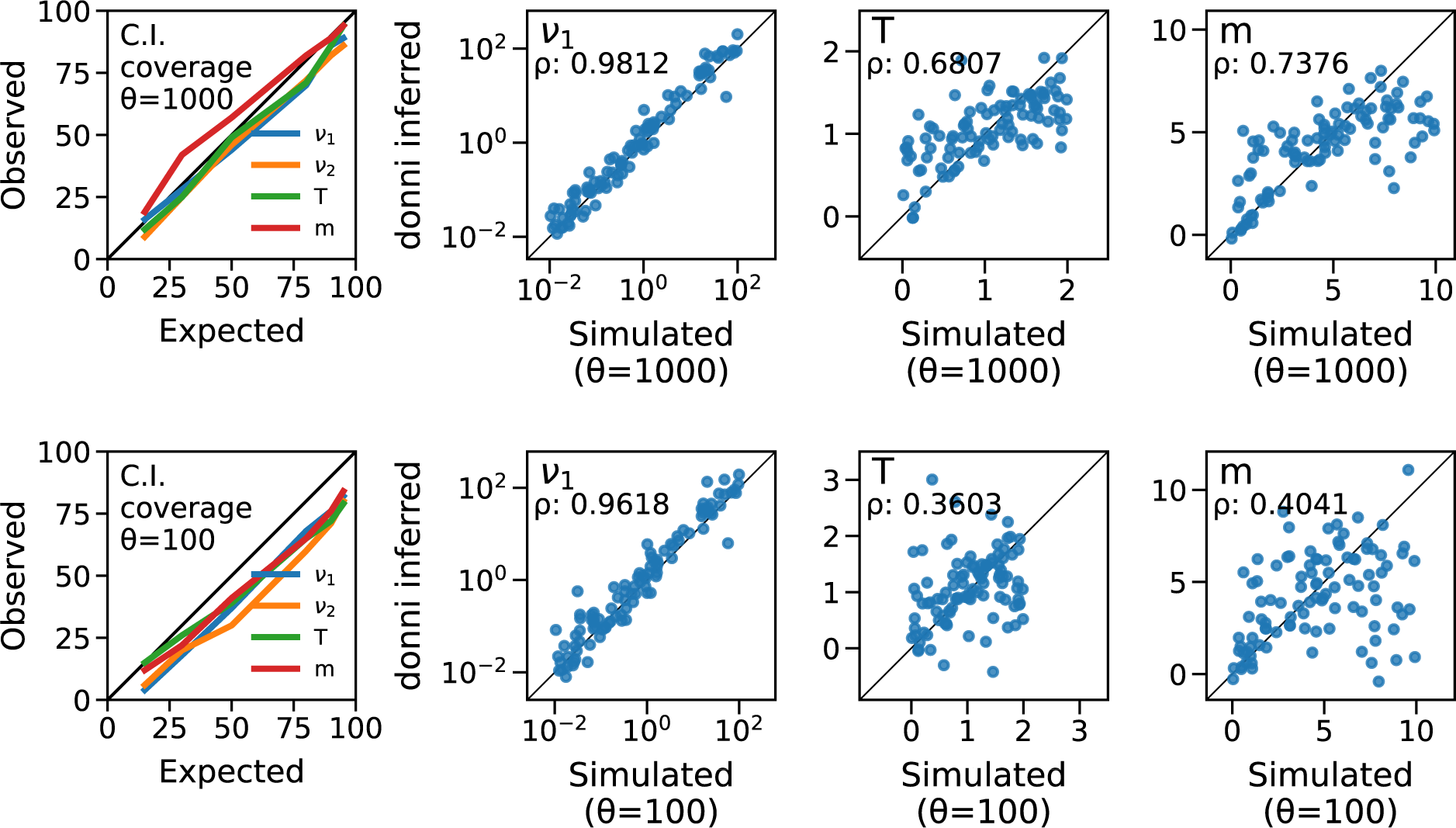
Inference accuracy and confidence interval calibration by donni on down-projected test AFS for the split-migration model. We simulated 100 test AFS with sample size 39 haplotypes per population then projected them to sample size 20 haplotypes per population. Top row is the result for test AFS projected from moderate variance (*θ* = 1000) AFS and bottom row is for test AFS projected from high variance (*θ* = 100) AFS.

**Figure S10:**
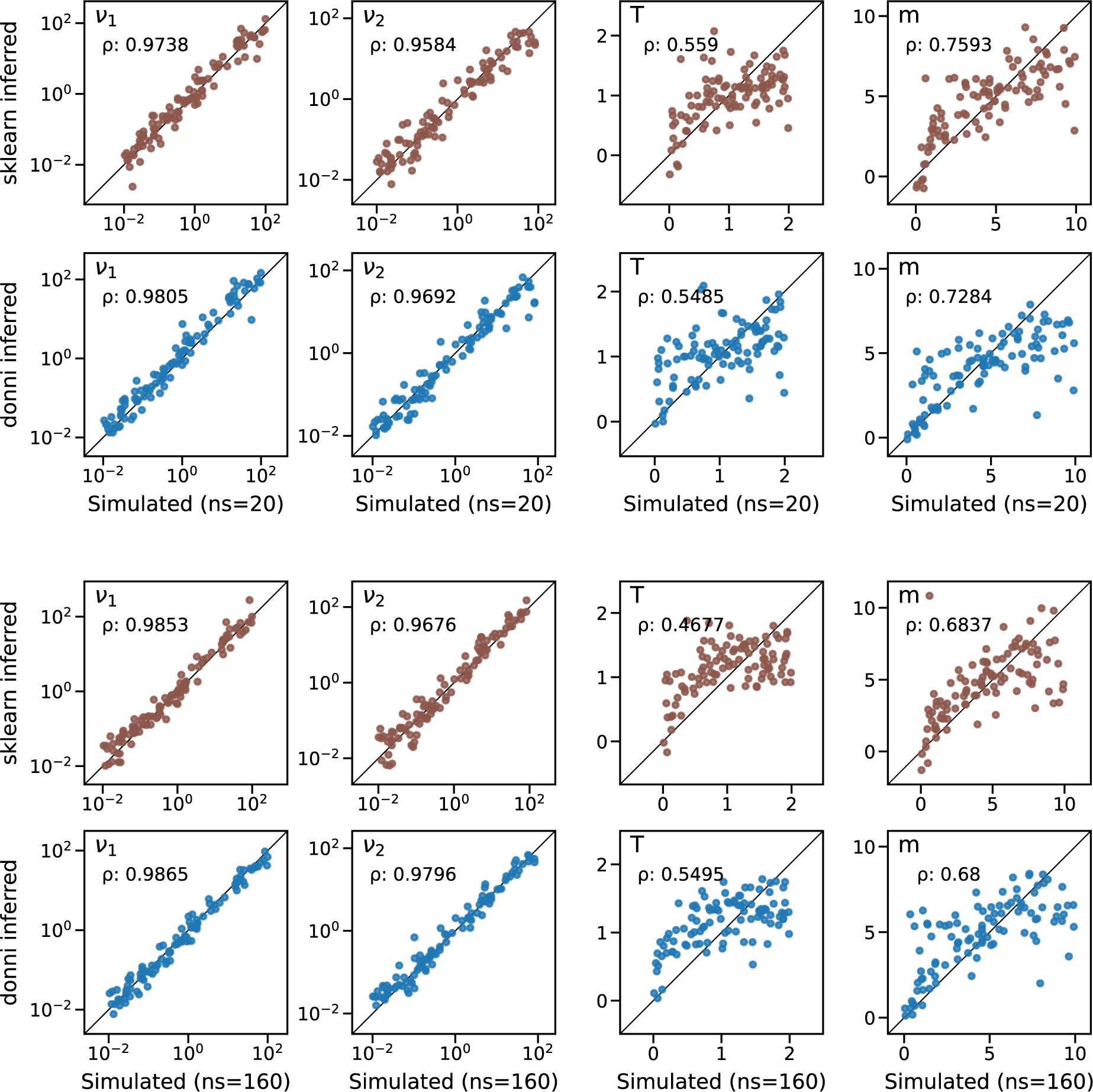
Inference accuracy by scikit-learn multi-output network compared with donni’s single-output network for the split-migration model with sample sizes 20 and 160 haplotypes per population. scikit-learn multi-output network is one network network trained to predict all parameters in a demographic model, whereas donni trains a single network for each parameter. We used the same test AFS simulated with moderate variance (*θ* = 1000) for sklearn and donni.

**Table S1:** donni inference accuracy on 1000 test AFS with moderate variance (*θ* = 1000) and the best hidden layer sizes (HLS) architecture for demographic history models in this study.

| Model | Parameter | RMSE | Spearman's $\rho$ | network architecture (HLS) |
| --- | --- | --- | --- | --- |
| 1-population | $\nu$ | 201.9 | 0.714 | (64, 12) |
| Two epoch | $T$ | 0.72 | 0.514 | (48, 9) |
| ($ns = 20$ haplotypes) | $misid$ | 0.037 | 0.905 | (64, 12) |
| 2-population | $\nu_1$ | 16.223 | 0.981 | (48, 12) |
| Split Migration | $\nu_2$ | 10.713 | 0.977 | (64, 12) |
| ($ns = 20$ haplotypes | $T$ | 0.444 | 0.65 | (32, 16) |
| per population) | $m$ | 2.008 | 0.714 | (48, 12) |
| | $misid$ | 0.019 | 0.968 | (48, 8) |
| 2-population | $\nu_1$ | 7.616 | 0.987 | (64, 16) |
| Split Migration | $\nu_2$ | 5.828 | 0.985 | (48, 12) |
| ($ns = 160$ haplotypes | $T$ | 0.455 | 0.609 | (48, 8) |
| per population) | $m$ | 1.927 | 0.736 | (64, 16) |
| | $misid$ | 0.012 | 0.987 | (64, 8) |
| 3-population | $\nu_{Af}$ | 11.504 | 0.985 | (64, 16) |
| Out of Africa | $\nu_B$ | 23.7371 | 0.495 | (64, 16) |
| ($ns = 20$ haplotypes | $\nu_{Eu0}$ | 22.609 | 0.451 | (48, 8) |
| per population) | $\nu_{Eu}$ | 17.67 | 0.921 | (48, 16) |
| | $\nu_{As0}$ | 24.06 | 0.405 | (64, 16) |
| | $\nu_{As}$ | 14.435 | 0.934 | (64, 16) |
| | $T_{Af}$ | 0.299 | 0.435 | (64, 4) |
| | $T_B$ | 0.322 | 0.366 | (48, 12) |
| | $T_{EuAs}$ | 0.303 | 0.515 | (64, 12) |
| | $m_{AfB}$ | 2.919 | 0.092 | (64, 16) |
| | $m_{AfEu}$ | 2.038 | 0.733 | (16, 16) |
| | $m_{AfAs}$ | 2.075 | 0.712 | (64, 8) |
| | $m_{EuAs}$ | 2.171 | 0.676 | (64, 12) |
| | $misid$ | 0.014 | 0.987 | (48, 16) |

**Table S2:** dadi demographic parameter range used for simulation in this study.

| Parameter | Symbol | Lower bound | Upper bound |
| --- | --- | --- | --- |
| Population size change | $\nu$ | 0.01 | 100 |
| Time of event(*) | $T$ | 0.01 | 2 |
| Migration rate | $m$ | 0 | 10 |
| Ancestral state misidentification | $misid$ | 0 | 0.25 |

\* For models with more than one $T$ parameter, the range specified for time $T$ applies to the sum of all $T$ parameters ( $T_{sum}$ ). For each demographic model, we drew different $T_{sum}$ values according to the desired number of data sets. For each data set, we then drew a set of $T$ parameters that sum to $T_{sum}$ by sampling from the Dirichlet distribution.

**Table S3:** donni inferred compared to dadi inferred parameter values in genetic units for the Out-of-Africa model using data from (Gutenkunst et al. 2009).

| Parameter | dadi | donni | donni 95% C.I. |
| --- | --- | --- | --- |
| $\theta$ | 2788.2 | 2644.9* | n.a. |
| $\nu_{Af}$ | 1.68 | 2.029 | 0.877 - 4.695 |
| $\nu_B$ | 0.287 | 0.192 | 0.011 - 3.500 |
| $\nu_{Eu0}$ | 0.129 | 0.145 | 0.004 - 5.627 |
| $\nu_{Eu}$ | 3.74 | 1.068 | 0.149 - 7.658 |
| $\nu_{As0}$ | 0.070 | 0.045 | 0.001 - 2.433 |
| $\nu_{As}$ | 7.29 | 1.276 | 0.200 - 8.135 |
| $m_{AfB}$ | 3.65 | 5.089 | -0.378 - 10.556 |
| $m_{AfEu}$ | 0.44 | 1.673 | -1.082 - 4.428 |
| $m_{AfAs}$ | 0.28 | 0.373 | -1.591 - 2.337 |
| $m_{EuAs}$ | 1.40 | 4.871 | -0.262 - 10.004 |
| $T_{Af}$ | 0.607 | 0.432 | -0.124 - 0.989 |
| $T_B$ | 0.396 | 0.22 | -0.310 - 0.749 |
| $T_{EuAs}$ | 0.058 | 0.119 | -0.084 - 0.321 |

\* donni infers a slightly negative value for the probability of ancestral state misidentification ( $misid = -0.00048$ ). These data were previously corrected for ancestral state misidentification using the approach of Hernandez et al. (2007). We thus rounded to $misid = 0$ when calculating $\theta$ .

**Table S4:** Hyperparmeters tuned with KerasTuner for each demographic model parameter.

| Tuner | Hyperparameter | Value range |
| --- | --- | --- |
| Hyperband | First hidden layer | 16, 32, 48, 64 |
|  | Second hidden layer | 4, 8, 12, 16 |
|  | Learning rate | log sampling, [0.0001, 0.01] |
|  | Max epochs | 100 |
| RandomSearch | Learning rate | log sampling, [0.0001, 0.01] |
|  | Max trials | 100 |

## Notes

https://github.com/lntran26/donni

### Competing Interest Statement

The authors have declared no competing interest.

### Summary of Updates

Core of inference engine has been changed to Mean Variance Estimation neural networks. Other revisions throughout.

